# INO80 Chromatin Remodelling Coordinates Metabolic Homeostasis with Cell Division

**DOI:** 10.1101/169128

**Authors:** Graeme J. Gowans, Alicia N. Schep, Ka Man Wong, Devin A. King, William J. Greenleaf, Ashby J. Morrison

## Abstract

Adaptive survival requires the coordination of nutrient availability with expenditure of cellular resources. For example, in nutrient-limited environments, 50% of all *S*. *cerevisiae* genes synchronize and exhibit periodic bursts of expression in coordination with respiration and cell division in the Yeast Metabolic Cycle (YMC). Despite the importance of metabolic and proliferative synchrony, the majority of YMC regulators are currently unknown. Here we demonstrate that the INO80 chromatin-remodelling complex is required to coordinate respiration and cell division with periodic gene expression. Specifically, INO80 mutants have severe defects in oxygen consumption and promiscuous cell division that is no longer coupled with metabolic status. In mutant cells, chromatin accessibility of periodic genes, including TORC-responsive genes, is relatively static, concomitant with severely attenuated gene expression. Collectively, these results reveal that the INO80 complex mediates metabolic signaling to chromatin in order to restrict proliferation to metabolically optimal states.

## INTRODUCTION

The coordination of cellular function with the environment is essential for adaptation and survival. Cells and organisms have a remarkable ability to sense diverse (i.e. nutrient-rich or -limiting) environments and reprogram their energy metabolism and proliferative capacity accordingly. Limiting nutrient environments are ubiquitous throughout nature and range from competitive microorganism growth environments to niches surrounding developing tissues in multicellular organisms. Failure to adapt can lead to cell death, developmental defects, and disease. Indeed, energy metabolism alterations are a major contributing factor for many pathologies, including cancer, cardiovascular disease, and diabetes, which together account for two-thirds of all deaths in most industrialized societies.

Adaptive cellular responses are often achieved by rapid inducible changes in gene expression^1^. For example, metabolic adaptation in the budding yeast *Saccharomyces cerevisiae* is achieved, in part, by coordinated regulation of gene expression, cell division, and metabolic status of the cell in low nutrient environments. This coordination can be observed when yeast are grown in low glucose conditions that mimic natural environments in the wild. In such glucose-deprived conditions, yeast quickly synchronize their metabolic processes and undergo coordinated and periodic bursts of respiration during a phenomenon known as the Yeast Metabolic Cycle (YMC)^2,3^.

Previous examination of gene expression in the YMC has revealed that over half of all transcripts undergo periodic expression, with peak levels occurring in one of three phases of the YMC (Fig. 1a). For example, genes involved in protein synthesis are induced while oxygen is rapidly consumed during the Oxidative phase (OX). Following OX, when oxygen consumption is reduced (Reductive Building phase, RB), genes involved in DNA replication and mitochondrial biogenesis are up-regulated. Subsequently, genes involved in glycolysis and the environmental stress response are up-regulated during periods of low oxygen consumption in the Reductive Charging (RC) phase. This temporal organization facilitates ‘just in time’ coordination of cellular function, wherein genes are transcribed just prior to the utilization of their encoded proteins within a particular metabolic or cell cycle pathway^4–7^. Temporal compartmentalization of gene expression also conserves cellular energy because futile metabolic reactions are avoided. Furthermore, previous research has found that periodically expressed genes in multiple species are among the most energetically expensive to transcribe and translate^4,8^. As such, coordinated periodic transcription -- rather than constitutive expression - maximizes the efficiency of limited cellular resources.

**Figure 1.**
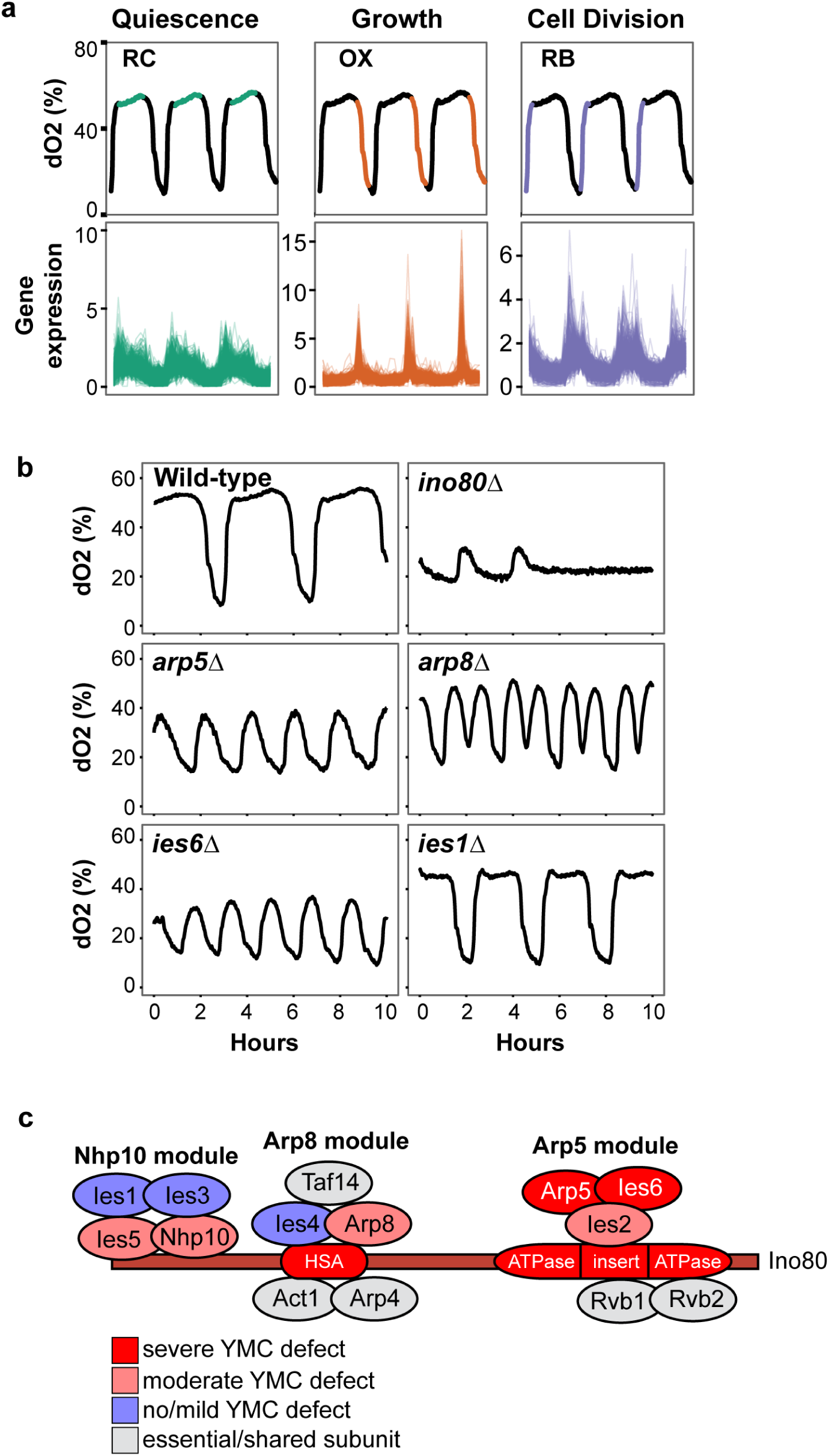
The INO80 complex is essential for respiration oscillations. **(a)**Organization of the yeast metabolic cycle (YMC). Top panels, three respiration cycles are shown with corresponding percentage of dissolved oxygen (dO2) in culture. Bottom panels, relative fold change gene expression of previously classified periodic genes^3^. RC is reductive charging (1508 periodic genes), OX is oxidative (1016 periodic genes), RB is reductive building (975 periodic genes). **(b)** YMC respiration (dO2) traces in wild-type and indicated mutant strains lacking subunits of the INO80 complex over a 10-hour period. **(c)** Illustration of the INO80 complex with structural modules noted from^27^. Colours denote the severity of YMC defects observed upon deletion of indicated subunit.

Importantly, periodic gene expression is an evolutionarily-conserved process. These highly robust gene expression cycles are observed in both lab and wild prototrophic yeast strains, and in single cells from asynchronous populations^9–11^. Oscillating gene expression has also been observed in nearly one-fifth of all genes in the developing *C*. *elegans* larvae^12^. Similarly, mammals undergo circadian rhythms, with oscillations in gene transcription and metabolic pathways coordinated with the day/night cycle^13^. Analysis of the mouse transcriptome across multiple tissues revealed that 43% of protein coding genes exhibit circadian-driven expression oscillations^14^.

Tight regulation of periodic gene expression not only promotes metabolic efficiency, as previously mentioned, but also optimizes cell growth and division. This is evident by the temporal constraint of cell division in both the circadian cycle and YMC ^2,3,11,15,16^. In the YMC specifically, cell division is gated within the RB phase, purportedly to shield replicating DNA from genotoxic reactive oxygen species^17^. Cell division frequency can be dynamically altered by nutrient availability and is absent without metabolic oscillations^18,19^. These studies highlight the critical importance of coordinating cell growth and division with metabolic environments. Without this synchrony, viability of the entire population can be jeopardized as additional cells increase competition for limiting nutrients.

Although the importance of synchronous metabolic and cell division oscillations is becoming increasingly clear, the identification of regulatory mechanisms for these processes is currently lacking. For example, the co-expression of periodic genes in the YMC has been proposed to be under the control of shared transcriptional regulators that exhibit periodic expression themselves. Although a few of these transcription factors have been identified^20^, the vast majority of YMC transcriptional regulation currently remains unexplained. However, an emerging concept in the field of cellular metabolism is that chromatin modification is an ideal mechanism to orchestrate gene expression in coordination with the metabolic environment^21^. This is primarily because many chromatin modifiers are abundant, regulate a large number of genes, and require intermediary metabolites as enzymatic cofactors. Thus, they can “communicate” metabolic status to the global chromatin environment. Moreover, epigenetic alterations can be rapidly and reversibly induced, and are capable of dynamically directing gene expression in coordination with changing nutrient environments.

We have recently identified the evolutionarily conserved INO80 complex as a regulator of metabolic function^22^. The INO80 chromatin-remodeling complex restructures and repositions nucleosomes *in vitro*^23,24^. *In vivo*, INO80 is enriched at the +1 nucleosome that plays critical roles in defining chromatin accessibility at promoters^22,25^. The *S*. *cerevisiae* INO80 chromatin remodeller is composed of 15 subunits^26^ that constitute four structurally distinct subunit modules along the Ino80 ATPase^27,28^. We found that the Arp5-Ies6 module, which is needed for chromatin remodelling catalytic activity^27–30^, regulates expression of genes in energy metabolism pathways. Specifically, *arp5Δ* mutants display downregulation of glycolytic genes and an upregulation of genes involved in the oxidative phosphorylation pathway^22^. Accordingly, mitochondrial potential and oxygen consumption are altered in *arp5Δ*, *ies6Δ*, and *ino80Δ* mutants.

To further investigate the relationship between INO80 and metabolic homeostasis, we examined the effect of INO80 deletion on metabolic and cell division oscillations in the YMC. In this report, we show that deletion of INO80 subunits severely disrupts metabolic cycling. RNA-seq and ATAC-seq analyses reveal global deviations in the expression of periodic genes in mutant cells, as well as alterations in chromatin accessibility across metabolic transcription factor motifs. Most notably, transcriptional effectors of the TOR pathway, a critical nutrient-sensing signaling pathway^31^, exhibit chromatin accessibility defects within their respective motifs. Furthermore, we demonstrate that TOR signaling and downstream INO80-regulated gene activity is critical for maintenance of the YMC. Finally, we show that disruption of the INO80 complex decouples cell division from metabolic oscillations. Collectively, our results establish the INO80 complex as a key regulator of metabolism that integrates nutrient signaling with gene expression and cell growth.

## RESULTS

### The INO80 complex is essential for respiration oscillations

To determine the role of INO80 in the YMC, we analyzed the effect of genetic disruption of each of the unique, non-essential subunits of the INO80 complex on YMC respiratory oscillations. We found that deletion of the Ino80 ATPase subunit had the most severe effect on the YMC organization, as the culture rapidly lost the ability to periodically respire (Fig. 1b). Deletion of *ARP8* resulted in a metabolic cycle that prematurely entered the OX phase at the midpoint of RC. While not as severe as the *ino80*Δ mutant, deletion of either *ARP5* or *IES6* resulted in rapid oscillations in oxygen consumption (Fig. 1b), with each cycle lasting approximately half the length of a wild-type cycle. The *arp5Δ* and *ies6*Δ mutants exhibited similar phenotypes in the YMC, consistent with their physical association as a subcomplex both within and outside the INO80 complex ^22,27,28,30^. The difference in YMC length between *arp5Δ* and wild-type strains was largely contributed by a drastically reduced RC (low oxygen consumption) phase. A similar YMC altered profile was observed for the *ino80*Δ mutant before the cycles ceased (Fig. 1b).

Deletion of *IES5* or *NHP10* also disrupted the periodicity of the YMC (Supplementary Fig. 1a). Ies5 is part of the Nhp10 module^27^ and purification of the INO80 complex from an *ies5*Δ strain showed a dramatic reduction in Ies1, 3 and Nhp10 (Supplementary Fig. 1b), corroborating recent results^32^. Interestingly, deletion of *NHP10* and *IES5* do not result in fitness defects in rich media^33^ (Morrison lab, unpublished), while deletion of the catalytic subunits of the INO80 complex, such as *ARP8*, *ARP5* and *IES6*, reduces fitness^22^. Nevertheless, *nhp10*Δ and *ies5*Δ mutants both displayed defects in respiration cycles in the YMC. Thus, defects observed in the YMC do not always translate to fitness defects in unsychronized cells.

Deletion of other subunits of the INO80 complex, including *IES1*, *IES3* and *IES4* had only minimal effects on the YMC. A summary of the phenotypes within each structural module is illustrated in Fig. 1c. Purification of Ino80-Flag throughout the YMC from wild-type cells demonstrated that the complex composition is not dramatically altered during the respiration oscillations (data not shown); thus, reconfiguration of the complex does not regulate normal YMC kinetics. Notably, the YMC defects observed in mutants of the INO80 complex are not common to all remodelers, as deletion of *SWR1*, a subunit of the SWR1 chromatin-remodeling complex and another chromatin remodeler in the INO80 subfamily^34^, had no effect on the cycle (Supplementary Fig. 1c).

### Disruption of the INO80 complex alters global transcription across the YMC

In asynchronous cultures, the INO80 complex regulates the expression of an abundance of genes enriched in metabolic processes^22^. To assess differences in transcript levels across the YMC, we performed RNA-seq analysis of both wild-type and *arp5Δ* cultures at six different time points during the cycle. We selected the *arp5Δ* mutant as it has a severely disrupted profile, whilst still maintaining sufficient cycles for sampling. To examine the relationship between samples, principal component analysis (PCA), a statistical procedure that reduces the dimensionality and complexity in large data sets^35^, was performed. In this analysis, samples that are most similar are clustered closer together. Principal components 1 and 2 revealed that wild-type samples (triangles) are roughly arranged in a circle, mirroring the cyclical nature of the system (Fig. 2a). A similar pattern is observed for the *arp5Δ* data (circles), although the samples are noticeably clustered closer together, indicating that the differences between these samples are smaller than the differences observed between the wild-type samples.

**Figure 2.**
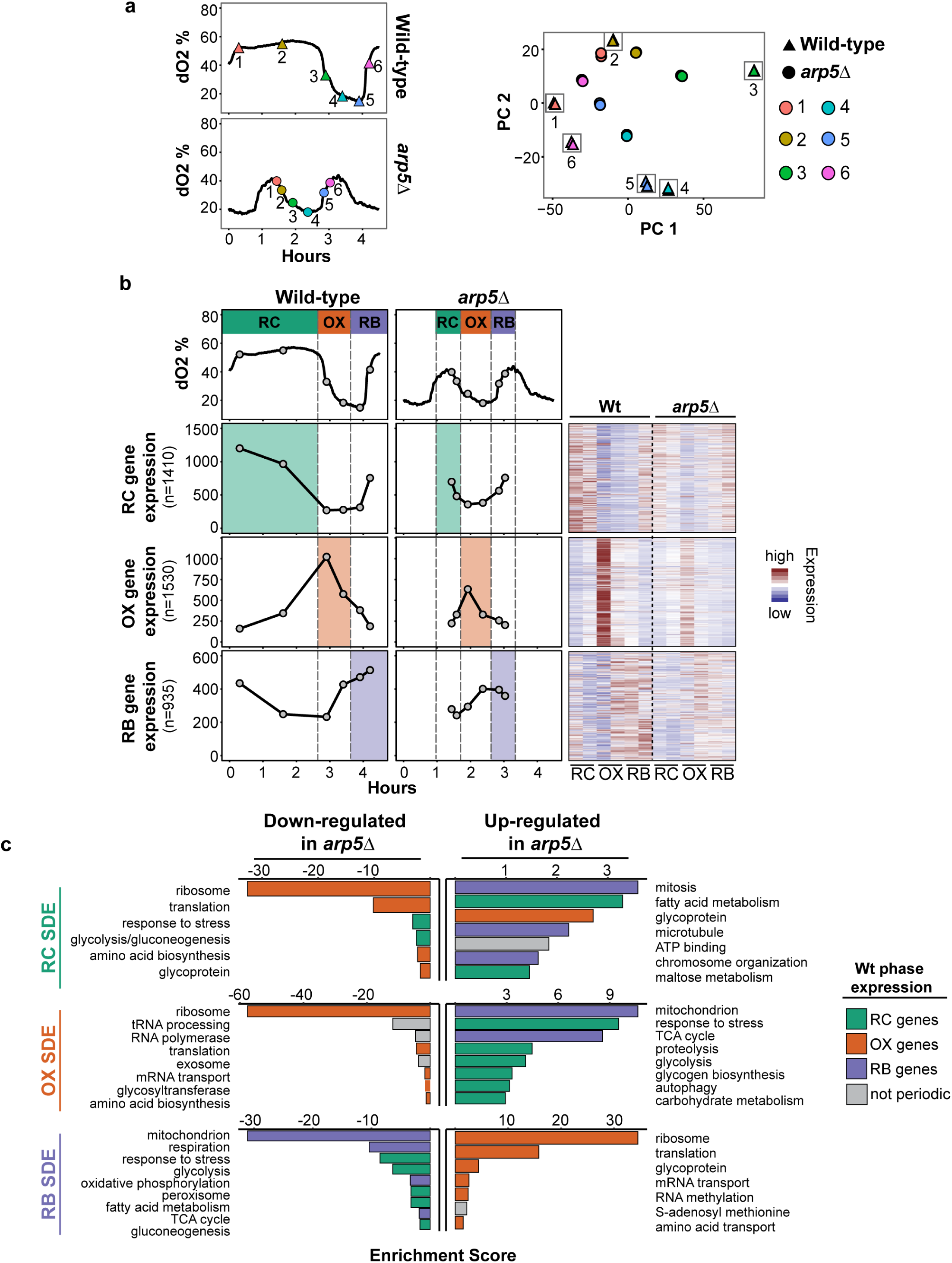
Disruption of the INO80 complex alters global transcription across the YMC. **(a)**Left panels, YMC dissolved oxygen (dO2) traces in wild-type and *arp5Δ*mutant. Samples were taken at indicated time-points (coded by colour) for RNA-seq and principal component analysis. Right panels, principal components (PC) 1 and 2 are shown. Technical replicates are displayed for each time-point and separated by strain. **(b)** Left panels, mean expression of periodic genes in wild-type (Wt) and *arp5Δ* strains within each YMC phase. n= number of periodic genes in each phase previously classified^36^. Expression of each gene is Fragments Per Kilobase of transcript per Million mapped reads (FPKM). Right panels, expression of each individual gene is shown as heatmaps (scaled by row). **(c)** DAVID functional enrichment analysis of significantly differentially expressed (SDE) genes at comparable time-points between wild-type and *arp5Δ* mutant (refer to Methods for more detail). Bar colours denote the phase in which periodic genes are associated with indicated functional annotation in the wild-type strain. RC is reductive charging, OX is oxidative, RB is reductive building. Enrichment scores are -log10(p-value).

To determine if specific metabolic pathways were mis-regulated in the *arp5* mutant, we next compared the expression of previously characterized periodic genes^36^ within each phase of the YMC (Fig. 2b). In the *arp5Δ* strain, although transcripts peaked during the expected phases, expression was markedly attenuated compared with wild-type. Significant differentially expressed (SDE) gene analysis between the mutant and wild-type during each phase of the YMC supported the observation of “muted” cycles in the *arp5Δ* mutant (Fig. 2c and Supplementary table 1). For example, pathways with peak expression in the wild-type OX phase, such as ribosome biogenesis, are also expressed in *arp5Δ* mutant at the same time-point. However, these transcripts are identified as significantly down-regulated when compared to wild-type cells because peak expression is dampened in mutant cells (Fig. 2c). Conversely, many of these OX phase pathways are comparatively up-regulated in the *arp5Δ* mutant during the RB phase, because they are not repressed to the same degree as in wild-type cells. This data demonstrates that in the *arp5Δ* mutant, periodic genes are generally expressed in the corresponding phase. However, the amplitude of periodic expression is dramatically decreased, resulting in YMC phasing that is less defined.

### Cell division is disconnected from the YMC in the *arp5*Δ mutant

Additional analysis was then performed to identify periodic genes in the *arp5Δ* mutant that do not exhibit expression in the expected YMC phase. Unsupervised clustering (*k*-means) analysis was performed on previously defined periodic genes^36^ after normalizing (z-score) by transcript, time-point, and strain, thereby adjusting the muted expression defects observed in *arp5Δ* cells. For both wild-type and *arp5Δ* cells, the vast majority of periodic genes (>3800 genes) clustered into one of three distinct patterns that reflect RC, OX, and RB peak mean expression (Fig. 3a and Supplemental table 2). This supports the observation that while gene expression is dramatically muted in *arp5Δ* mutants, periodic expression of many genes is still detectable. However, we identified 722 genes in *arp5Δ* mutants with peak expression and *k*-means clustering in a different phase than expected, representing genes with severe defects in periodic transcriptional regulation (Supplemental table 2). Functional annotation analysis demonstrates that these genes are significantly enriched in cell cycle and mitotic pathways (Fig. 3b). As previously discussed, cell division and DNA replication are tightly coupled with the RB phase of the YMC^2,3,11,17^. Additionally, genes involved in mitosis and chromosomal segregation were significantly up-regulated in the RC phase of *arp5Δ* cells (Fig. 2c). Thus, cell cycle gene expression is dramatically altered in the *arp5Δ* mutant YMC.

**Figure 3.**
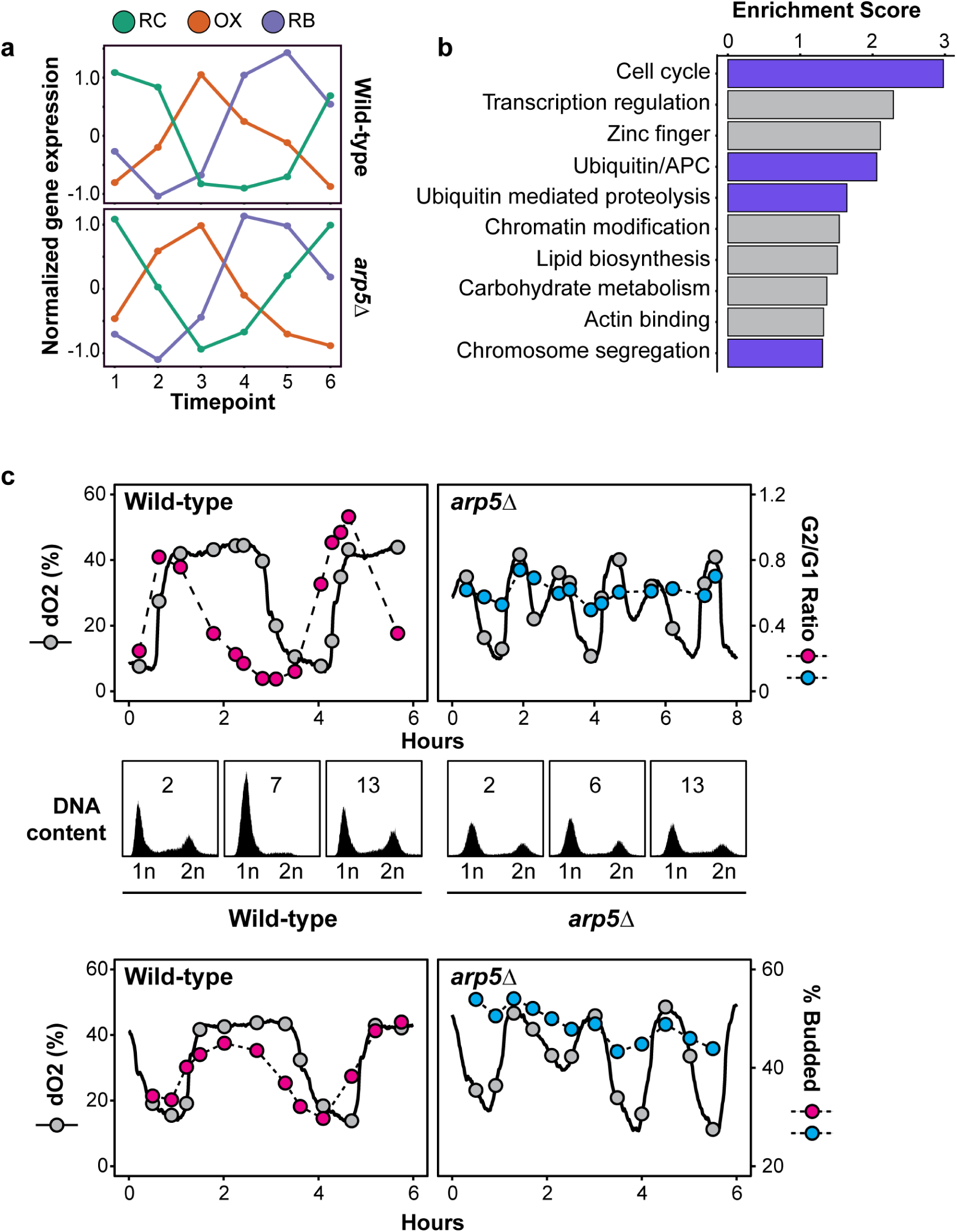
Cell division is disconnected from the YMC in the *arp5Δ* mutant. **(a)** *k*-means clustering of previously classified periodic genes^36^ following z-score normalization of RNA-seq data within each gene, time-point, and strain. RC is reductive charging, OX is oxidative, RB is reductive building. **(b)** DAVID functional enrichment of genes in the *arp5Δ* mutant that do not cluster in the corresponding YMC phase. Annotations related to cell cycle are highlighted. Anaphase promoting complex is APC. Enrichment scores are -log10(p-value). **(c)** *arp5Δ* mutants have defects in cell cycle synchronization across the YMC. Samples were taken at indicated time-points across wild-type and *arp5Δ* YMCs. Dissolved oxygen (dO2) traces for each strain are shown. DNA content was assessed by SYTOX Green staining followed by flow cytometry. The ratio of G2 to G1 cells and representative histograms from indicated samples are shown. **(d)** Analysis of budding cells across the YMC. Samples were taken across wild-type and *arp5Δ* YMCs and the number of budding cells, indicative of M phase entry, was assessed via microscope at each time-point.

To further investigate the disruption of cell cycle kinetics in *arp5Δ* cells, we monitored DNA content and cell budding across the YMC. As previously reported, the wild-type RB phase was tightly coupled to DNA replication (Fig. 3c) and cell division (Fig. 3d). However, in the *arp5Δ* mutant, DNA replication was uncoupled from the YMC, as we observed a population of cells with 2N DNA content and budding cells at all phases. The cell density of the *arp5Δ* culture remained constant and comparable to wild-type in the bioreactor with constant perfusion of media. Thus, these defects in the *arp5Δ* mutant are not due to cell cycle arrest or cell death. Conversely, these results suggest that disruption of the INO80 complex decouples cell division from the metabolic state of the cell.

### Loss of INO80 function alters global chromatin accessibility in the YMC

To assess the influence of the INO80 chromatin remodelling in the YMC, we utilized the Assay for Transposase-Accessible Chromatin using sequencing (ATAC-seq)^37^. ATAC-seq uses a hyperactive Tn5 transposase that preferentially inserts specific adaptors in regions of accessible chromatin. Samples were taken at the same time-points as the RNA-seq (Fig. 2a). Similar to the RNA-seq results, PCA illustrates the cyclical relationship among samples (Fig. 4a) within PC2 and PC3, while PC1 largely partitioned wild-type and the *arp5Δ* samples separately (Supplementary Fig. 2a). The distribution of samples in the PCA plot is indicative of periodic fluctuations in chromatin accessibility across the YMC. However, the *arp5Δ* samples were clustered in the center of the plot, suggesting that the differences between these samples were much less pronounced than in the wild-type (Fig. 4a).

**Figure 4.**
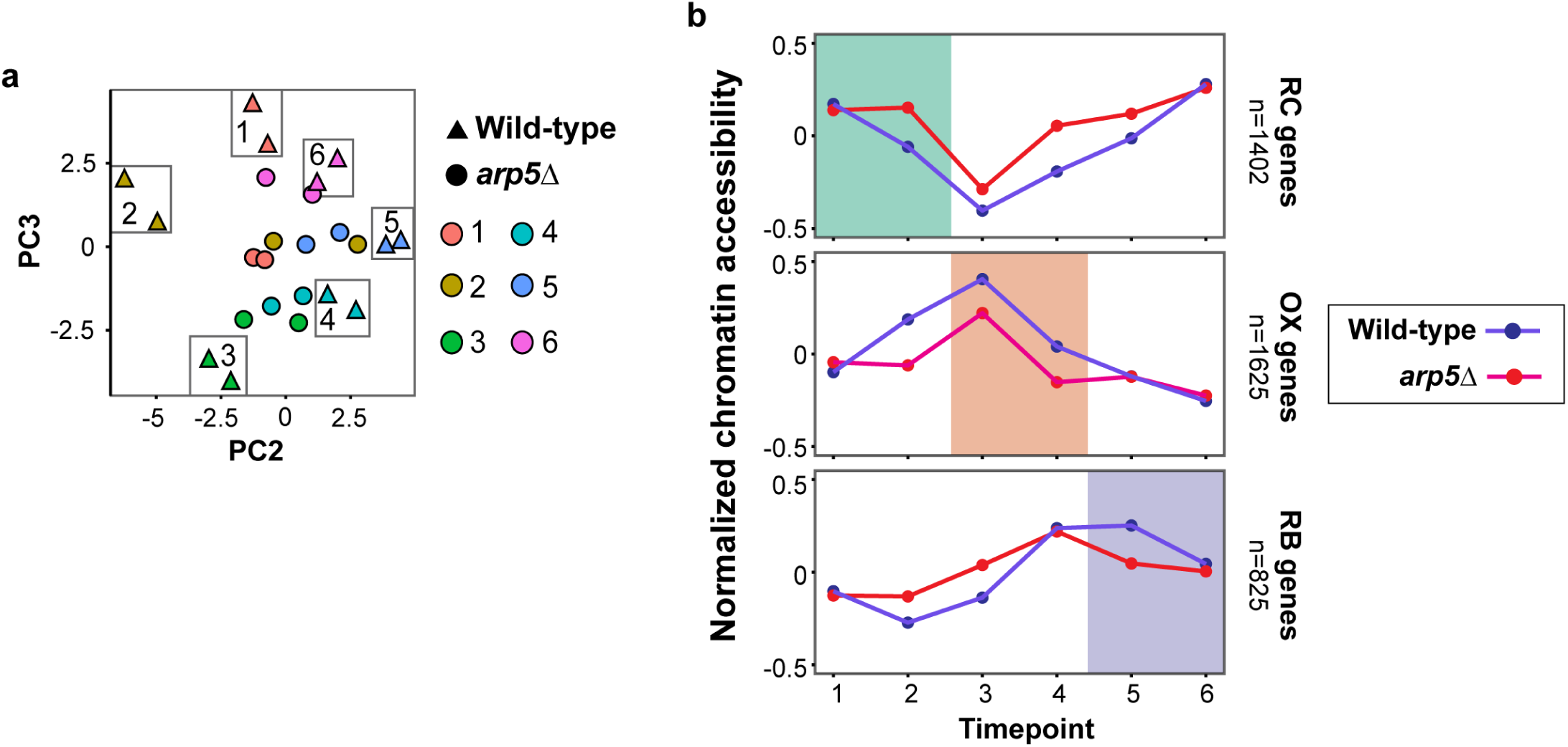
Loss of INO80 function alters global chromatin accessibility in the YMC. **(a)** Principal component analysis (PCA) plot of ATAC-seq data from samples taken at same time-points shown in Fig. 2a. PC2 and PC3 are shown. Replicates are displayed for each time point and strain (coloured as in Fig. 2a). **(b)** Comparison of mean ATAC-seq scores for periodic genes from each phase of wild-type and *arp5Δ* following z-score normalization within each gene and time-point. Reductive charging (RC), oxidative (OX), and reductive building (RB) phases are highlighted.

To explore potential mechanisms of INO80-mediated transcription regulation, we then investigated ATAC accessibility across all periodic gene promoters, defined as −400bp downstream and 100bp upstream of the transcriptional start site (TSS). ATAC scores largely correlated with gene expression (r >0.7), with increased accessibility corresponding with the phase of periodic expression (Fig. 4b). However, accessibility was more static in *arp5Δ* cells compared to wild-type. This data reveals that disruption of INO80 chromatin-remodeling results in global chromatin accessibility defects that reduce the plasticity of the chromatin template during metabolic oscillations.

### Metabolic gene promoters are dependent on INO80-facilitated chromatin accessibility

In order to identify individual transcription factors that are influenced by INO80 chromatin remodelling, chromatin accessibility was assessed for promoters of specific transcription factor (TF) motifs. The presence of each annotated transcription factor (n=177) was assessed for all promoters, defined as −400bp downstream and 100bp upstream relative to the TSS. Results revealed that, similar to the data in Figure 4b, the wild-type data was much more dynamic than the mutant (Fig. 5a). In particular, several motifs exhibited comparably large fluctuations in accessibility (Fig. 5b), suggesting a relatively large dependence on chromatin manipulation for regulation of those genes. Indeed, periodic expression of these highlighted TF-regulated genes is largely coordinated with accessibility in wild-type cells (Fig 5c). However, in the *arp5Δ* mutant, these motifs lacked dynamic accessibility and exhibited significant differential accessibility between wild-type and *arp5Δ* cells (Benjamini-Hochberg adjusted p<0.01) (Fig. 5b). In addition, corresponding gene expression is deregulated (Fig. 5c).

**Figure 5.**
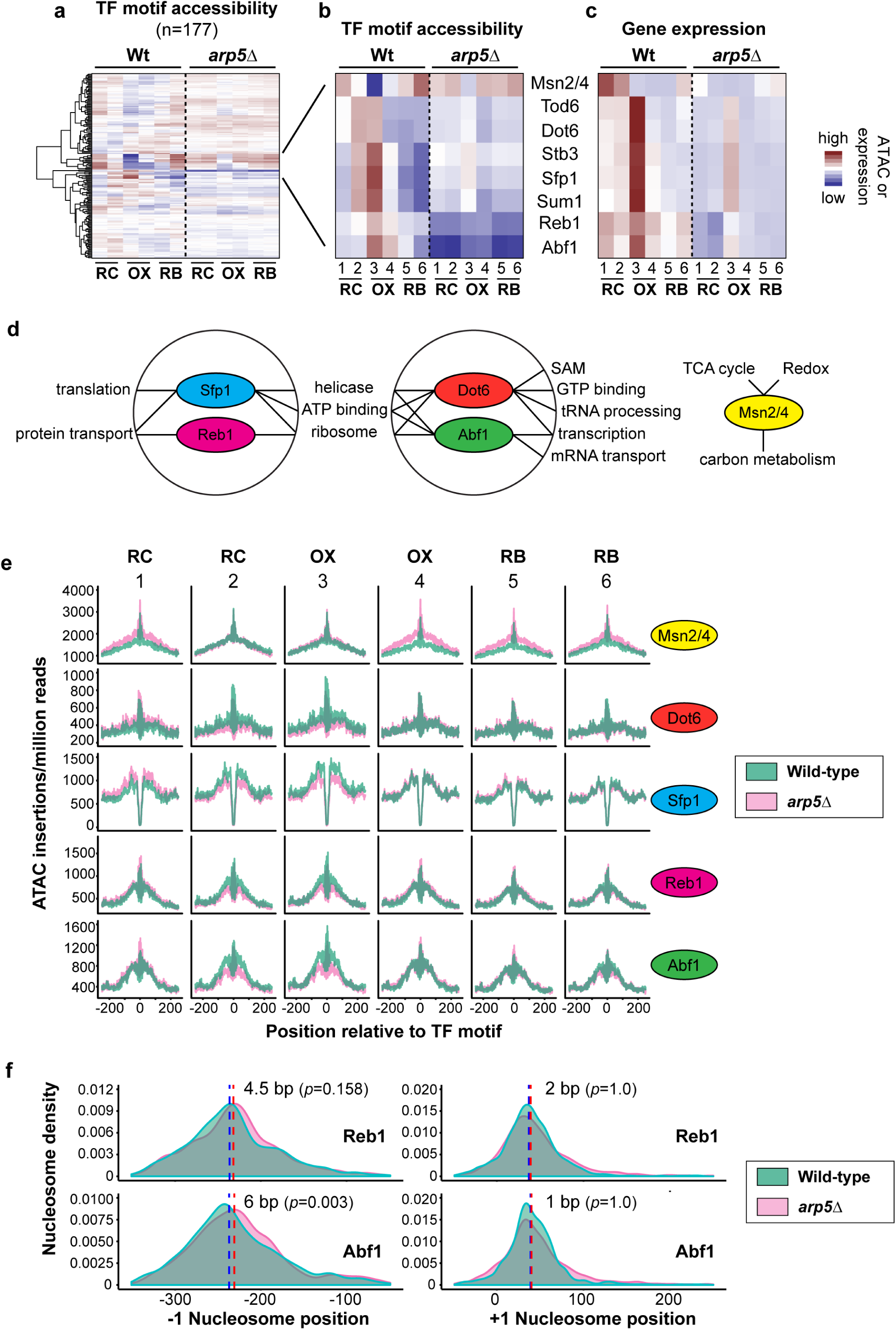
Metabolic gene promoters are dependent on INO80-regulated chromatin accessibility. **(a)** ATAC accessibility scores for promoters (400 bp upstream to 100 bp downstream of TSS) containing each motif were determined using ChromVar^70^ and organized by the presence of transcription factor (TF) motifs. **(b)** Magnification of indicated region in (a). TF motif in indicated promoter is shown. **(c)** Mean RNA-seq Fragments Per Kilobase of transcript per Million mapped reads (FPKM) for genes that contain indicated transcription factor motifs in their promoters. **(d)** DAVID functional enrichment of genes containing indicated transcription factor motifs. Only significant enrichments (enrichment score>1.3 (-log 10 p-value)) are shown. S-adenosyl-methionine is SAM, TCA cycle is tricarboxylic acid cycle. **(e)** ATAC accessibility is shown as the number of Tn5 insertions per million reads around the indicated transcription factor motif for wild-type (green) and *arp5Δ* (pink) samples. Timepoints and YMC phase from Fig. 2a are shown at top. **(f)** +1 and −1 nucleosome densities, relative to transcription start sites containing Abf1 or Reb1 motifs, were obtained from JASPAR database^48^ and identified as previously described using NucleoATAC^70^. Wild-type (green) and *arp5Δ* (pink) samples across all time-points of the YMC are shown. Red/blue dotted lines denote the median nucleosome position for wild-type and *arp5Δ* respectively. p = 0.00260 (Abf1) and 0.15808 (Reb1) based on a permutation test and adjusting for multiple hypothesis testing.

Notably, these highlighted TF motifs are enriched in components of the energy-responsive TORC1 pathway (Fig. 5d). For example, in nutrient-rich growth conditions, Msn2/4 are inhibited by TORC1 via phosphorylation that results in their exclusion from the nucleus^38^. In nutrient-depleted growth conditions, these transcription factors translocate to the nucleus and express genes in the general stress response^39–41^. Dot6, Tod6 and Stb3 are phosphorylated by Sch9 in a TORC1-dependent manner to promote expression of ribosomal protein (RP) and ribosomal biogenesis (Ribi) gene expression^42,43^ (Fig. 5d). TORC1 also directly phosphorylates the transcriptional activator Sfp1 to facilitate its binding to RP gene promoters and subsequent expression^44,45^.

Interestingly, Reb1 and Abf1, which are critical factors in controlling nucleosome positioning at promoters^46^, are found in the promoters of Ribi genes and genes involved in protein and mRNA transport (Fig. 5d). Consistent with our observations, a recent study identified Reb1 and Abf1 at Ribi promoters using chromatin immunoprecipitation (ChIP) *in vivo*^47^. In *arp5Δ* mutant cells, Reb1 and Abf1 motifs were the most significantly statically inaccessible among all motifs and across all phases of the YMC (Benjamini-Hochberg adjusted p<0.001). Comparatively, in wild-type cells, the Reb1 and Abf1 motifs exhibited coordinated oscillations of accessibility and gene expression in the YMC (Fig. 5b and c).

We then examined accessibility at high resolution across the motifs of these transcription factors (Fig. 5e and Supplementary Fig. 2b). We observed that there were striking phase-dependent differences between wild-type and *arp5Δ* mutant cells. Specifically, when comparing these two strains, Dot6, Sfp1, Reb1, and Abf1 motifs show the greatest difference in accessibility during the OX phase (Fig. 5e). In wild-type cells, this peak of accessibility correlates with the peak of expression of genes containing these motifs in their promoters (Fig. 5b and c), and likely stems from a shared transcriptional program involved in ribosome biogenesis and translation (Fig. 5d). Conversely, regions containing Msn2/4 motifs have reduced accessibility during the OX phase, with comparably smaller differences between the wild-type and the *arp5Δ* mutant (Fig. 5b). However, at higher resolution, strain-dependent differences are observed within Msn2/4 motifs during the RB and RC phase (Fig. 5e). During these phases of the YMC, Msn2/4-dependent gene expression is induced in the wild-type (Fig. 5c) and is enriched in energy metabolism pathways (Fig. 5d). These results suggest that INO80 activity is necessary to create dynamically accessible chromatin at the motifs of transcription factors important YMC regulation.

As previously stated, INO80 is a chromatin remodeler that is enriched at the +1 nucleosome^22,25^. Thus, we also assessed promoter proximal nucleosome positioning across the YMC in both wild-type and *arp5Δ* cells. However, our ability to confidently call nucleosomes was confounded in the *arp5Δ* mutants due to an overall decrease in predicted nucleosome spanning fragments compared to wild-type samples. The precise origin of this technical limitation is not known. However, it may be a consequence of increased +1 nucleosome “fuzziness”, i.e. deviation in mean nucleosome positioning of individual +1 nucleosomes in a cell population, which was previously observed in *ino80Δ and arp5Δ* mutants^22^. For +1 nucleosomes that could be confidently called, no statistically significant deviations were detected proximal to transcription factor motifs when comparing wild-type to *arp5Δ* mutants (Fig. 5f and data not shown).

However, we did observe a significant change in the −1 nucleosome position for genes containing either the Abf1 or Reb1 transcription factor motifs. We observed differences in the size and significance of the median shift in nucleosome position depending on the motif identification method used. Using the JASPAR database^48^, we observed median shifts of 6 and 4.5 bp in −1 nucleosome position proximal to Abf1 (p-value=0.026) and Reb1 (p-value=0.158) motifs, respectively (Fig. 5f). Using previously published ChIP analysis in the absence of chromatin crosslinking, called <u>o</u>ccupied <u>r</u>egions of <u>g</u>enomes from <u>a</u>ffinity-purified <u>n</u>aturally <u>i</u>solated <u>c</u>hromatin (ORGANIC)^49^, we observed median shifts in −1 nucleosome position of 3.5 and 4 bp proximal to Abf1 (p-value=0.012) and Reb1 (p-value=0.004) motifs, respectively (Supplementary Fig 2c). No other statistically significant deviations were observed for nucleosomes proximal to other transcription factor motifs (data not shown). This data supports a previously unrecognized role for the INO80 complex in −1 nucleosome positioning at specific metabolic promoters.

For most transcription factors, these defects in accessibility, gene regulation, and nucleosome positioning do not appear to be due to general gene expression defects of the transcription factors themselves (Supplementary Fig. 2d). While many of these transcription factors display periodic expression, which has been previously reported for Msn4^20^, the pattern of expression does not always directly correlate with the expression of other genes that have the motif. However, notable exceptions are Tod6 and Sfp1, which have patterns of transcription factor expression that closely mirror the expression of genes with corresponding motifs in their promoters (Fig. 5c and Supplementary Fig. 2d). Thus, a portion of the defects in metabolic gene expression and chromatin architecture observed in *arp5Δ* cells may originate with alterations of transcription factor expression itself. However, the INO80 chromatin remodeling also plays a major role in regulating the chromatin architecture at the recognition motifs of several metabolic transcription factors. The combined effect of INO80 loss of function is a dramatically unresponsive and static chromatin architecture that does not exhibit periodic regulation characteristics of the YMC.

### The INO80 complex is a key regulator of TORC1-responsive gene expression

Of all the transcription factors assessed, Msn2 and 4 had the largest ATAC-seq variability across the YMC of wild-type cells, which is altered in the *arp5Δ* mutant (Fig. 5b). Because Msn2 and 4 are regulated by the TORC1 pathway, we sought to further determine the role of TORC1 in the YMC. We first examined Msn2-mediated transcription in greater detail by restricting analysis to highly confident Msn2-regulated genes that have been validated by ChIP (n=88)^50^. Again, we found that Msn2-dependent genes were markedly reduced in the OX phase of the wild-type YMC (Fig. 6a), when accessibility of Msn2 motif-containing genes is lowest (Fig. 5b). In the *arp5Δ* mutant, we observed lower expression of these Msn2-induced genes in the RC phase of the cycle (Fig. 6a), with generally static accessibility patterns across the entire YMC at corresponding motifs (Fig. 5b).

**Figure 6.**
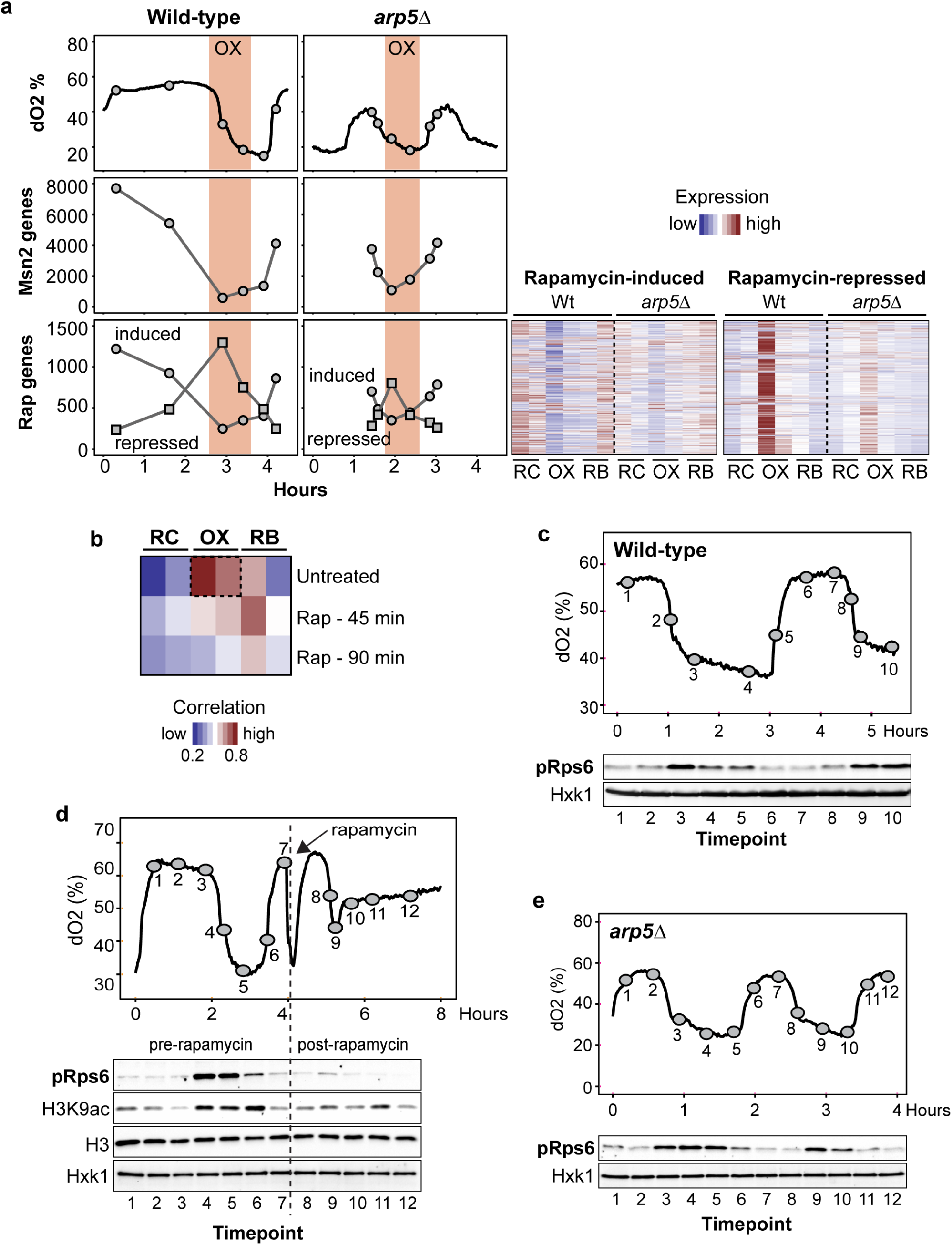
The INO80 complex is a key regulator of TORC1-responsive gene expression. **(a)** Top panel, same as Fig. 2a. YMC dissolved oxygen (dO2) traces in wild-type and *arp5Δ* mutant. RNA-seq samples were taken at indicated time-points. Mean expression is shown as Fragments Per Kilobase of transcript per Million mapped reads (FPKM) of Msn2-induced genes (middle panel, Msn2 genes) and rapamycin-sensitive genes (bottom panel. Rap genes) across the YMC. Right panels, gene expression heatmaps of rapamycin-sensitive genes scaled by row. **(b)** Correlation between the RNA transcript levels across the wild-type YMC and those in an asynchronous culture before and after rapamycin treatment for either 45 or 90 minutes (min). **(c)** TORC1 activity fluctuates across the YMC. Samples were taken across the wild-type YMC at the indicated time-points and used for western analysis with an antibody that recognizes the phosphorylated TORC1 substrate, mammalian homologue ribosomal protein S6 (Rps6). Hexokinase 1 (Hxk1) is shown as a loading control. **(d)** Rapamycin causes an arrest in the YMC. Samples were taken across a wild-type YMC at the indicated time-points before and after treatment with rapamycin (50 nM), indicated by dashed line. Western analysis was done as in Fig. 6c. **(e)** TORC1 signaling is periodic in the *arp5Δ* mutant. Samples were taken across the mutant YMC at the indicated time-points and used for western analysis as in Fig. 6c.

We then examined the requirement for INO80 on all TORC1-regulated gene expression by analyzing the expression of rapamycin-sensitive genes in the YMC. We identified both significantly up- (n=1177) and down-regulated (n=1249) genes from asynchronous cultures treated with rapamycin (Supplementary table 3). Analysis of these transcript levels showed that rapamycin-induced genes (TORC1-repressed) have reduced expression during the OX phase (Fig. 5A, bottom). The expression pattern of these TORC1-repressed genes closely mirrors the pattern of Msn2-regulatd genes. Conversely, rapamycin-repressed genes, (TORC1-induced), have peak expression during the OX phase. In the *arp5Δ* mutant, this divergent pattern between rapamycin-induced and -repressed expression was much less distinguished, demonstrating that TORC1-regulated gene expression is periodic in the YMC and dependent on INO80-mediated chromatin remodeling.

Overall, gene expression in untreated asynchronous cultures had the strongest correlation with the OX phase of the YMC, however rapamycin treatment greatly weakened this association (Fig. 6b). We then monitored TORC1 signaling by examining the phosphorylation of ribosomal protein S6 (Rps6)^51^, a downstream substrate in the TORC1 pathway^52^. Rps6 phosphorylation was nearly undetectable in the RC phase of the YMC and peaked during the OX phase of wild-type cells (Fig. 6c). These data reinforce the idea of the OX phase as being the main TOR-regulated growth phase of the YMC, with the other phases similar to quiescent or stationary phases in asynchronous cultures.

Due to the importance of TORC1-signaling in the YMC, treatment with rapamycin resulted in disruption of the YMC at the point of maximal TOR activity (Fig. 6d). Rapamycin prevented the increase in phospho-Rps6 and acetylation of a number of H3 lysine residues which are known to fluctuate across the YMC^53^ (Fig. 6d and data not shown). [Note that rapamycin was prepared in ethanol, which itself can be metabolized, accounting for the transient decrease in dO2. Addition of ethanol alone had no effect on subsequent cycles (Supplementary Fig. 3a)]. Addition of acetate has been shown to induce premature entry into OX phase^53^, and as we observed, an increase in TOR activity and histone acetylation (Supplementary Fig. 3b). Lastly, genetic deletion of *SCH9* or *RIM15*, downstream components of the TORC1 pathway^54,55^, also either prevented or disrupted metabolic cycles (Supplementary Fig. 3c).

As mentioned, TORC1-responsive gene expression is muted in *arp5Δ* cells (Fig. 6a). Interestingly, we observed that the induction of TORC1 signaling appeared normal in the *arp5Δ* strain, with a peak in the OX phase, despite the differences in YMC period. Thus, although the upstream TORC1 signaling is intact in the *arp5Δ* mutant, downstream TORC1-regulated gene expression is disrupted. These results demonstrate that the INO80 complex is a critical effector of TORC1-regulated gene expression that coordinates the quiescence (RC) to growth (OX) transition and maintains metabolic homeostasis.

## DISCUSSION

### The INO80 chromatin remodeler is a key regulator of metabolic function

Our data shows that the INO80 chromatin-remodeling complex plays a major role in maintaining metabolic homeostasis in yeast. We demonstrate that deleting the *ARP5* subunit of the INO80 complex causes differential expression of periodic genes and alterations in global chromatin architecture. The cumulative defect in *arp5Δ* mutant cells is a system level disorganization of the metabolic cycle, characterized by severely diminished ability to organize robust YMC phasing and coordinate cell division with metabolic timing.

The resulting effect of INO80 disruption is likely to contribute to metabolic inefficiency, whereby transcripts are produced out of synchrony with the function of the encoded protein. For example, in *arp5Δ* mutants, ribosomal gene expression is both produced (i.e. derepressed) out of phase and not fully activated during the optimal OX phase. Gene expression related to ribosome biogenesis involves nearly 10% of the genome, with many exhibiting >100 fold amplitude changes in the YMC^3^. It has been proposed that approximately 60% of total *S*. *cerevisiae* transcription is devoted to rRNA production and 50% of Pol II transcription is involved in ribosomal protein expression^56^. Thus, the process of translation creates a tremendous energy expenditure for the cell, which must be tightly regulated to ensure efficiency. In the YMC, gene expression programs related to translation peak during the OX phase, coincident with the production of ATP^57^ and acetyl-coA^53^ that can feed these energy demanding processes. Because mutants of the INO80 complex do not optimize this energy demanding process, all cellular processes that require coordination of function with energy availability are likely jeopardized.

Notably, we observe unrestricted cell division in *arp5*Δ mutants. The coordination of energy metabolism and cell division is vital to survival in competitive nutrient environments. Moreover, excessive proliferation that is disconnected from nutrient availability is characteristic of diseases such as cancer. For example, the mTOR signaling pathway is often constitutively active in cancer, promoting growth signaling and proliferation irrespective of metabolic environments^58^. In this study, we find that the INO80 complex is a critical component for enacting TOR responsive transcriptional programs. Interestingly, subunits of the evolutionarily-conserved INO80 complex are commonly amplified in many cancers, including 51% of lung squamous cell carcinoma^59^, 50% of pancreatic^60^, and 45% of bladder cancers^61^. Results from this study reveal the INO80 complex as a previously unrecognized regulator of proliferative capacity that restricts cell division to metabolically optimal states. Additional studies may further define the role of INO80 subunits in human metabolism and disease.

### The INO80 complex regulates chromatin architecture of metabolic genes

Our results also demonstrate that chromatin modification is an excellent means to regulate gene expression in coordination with changing metabolic environments. Previous research also supports this model by demonstrating that histone acetylation is tightly coupled to carbon availability^53^, which is converted to acetyl-CoA, an intermediary metabolite and cofactor for histone acetyltransferases.

We observe that chromatin accessibility is dynamically altered in the YMC. Specifically, metabolic promoters exhibit large fluctuations in accessibility that correspond to gene expression. For example, we observed that chromatin accessibility surrounding TORC1-responsive promoters fluctuated across the YMC and was largely static in the *arp5Δ* mutant. Thus, the organization of the YMC is particularly dependent on TORC1-regulated gene expression, which in turn is mediated by INO80 chromatin-remodeling.

We also observed dramatically reduced accessibility at Abf1 and Reb1 motif-containing promoters in the *arp5Δ* mutant. These promoters are found at ribosome biogenesis genes and, in wild-type cells, are among those with the largest fluctuations in accessibility across the YMC. Abf1 and Reb1 play major roles in establishing nucleosomal array positioning at promoters^46^. Abf1 motifs are enriched 250 bp upstream of the TSS^62^ and Reb1 localizes to the −1 nucleosome, near the nucleosome-free border upstream of the TSS^63^. We find that INO80 chromatin remodelling is needed to position the −1 nucleosome at promoters that contain Reb1 and Abf1 motifs. Interestingly, most of the −1 nucleosomes bound by Reb1 are found at divergently transcribed genes^63^, and disruption of Reb1 function causes a reduction in expression of these genes^64^. While loss of loss of Abf1 caused decreased nucleosome occupancy in upstream of the TSS^62^. Our data demonstrates that −1 nucleosome positioning by Abf1 and Reb1-mediated is dependent on INO80 chromatin remodelling, the disruption of which is detrimental for metabolic homeostasis.

It should be noted that some of the YMC defects observed in *arp5Δ* cells may be an indirect result of INO80 dysfunction. For example, *ARP5* deletion may directly result in a shortened RC phase, which consequently results in a relative elongation of the subsequent OX phase. In order to assess immediate effects of INO80 deletion, we attempted to use an auxin-based degron system to acutely disrupt the function of the INO80 complex^65^. However, the addition of the auxin chemical to a wild-type culture disrupted cycling, preventing the use of this system. Because the bioreactor system that controls the YMC is sensitive to exogenous perturbations, it is likely that inducible systems will be technically limited in the YMC. Nevertheless, the system is an excellent means to study cumulative metabolic effects resulting from constitutive genetic manipulations.

### The Yeast Metabolic Cycle is a powerful tool for studying metabolic signaling to chromatin

As mentioned, TORC1 signaling peaks during the transcriptional transition from quiescence (RC) to growth (OX), and rapamycin treatment inhibits this progression. It has also been reported that PKA-activity fluctuates across the YMC, peaking in OX phase, and that this was sensitive, in asynchronous cultures, to rapamycin treatment^53^. Interestingly, TOR and PKA have been shown to both regulate a number of transcription factors, including Dot6 and Tod6^43,66^. Thus, INO80 function may be critical for mediating the gene expression of multiple metabolic signaling pathways.

Much like the checkpoints that regulate the cell cycle, the YMC may similarly be regulated by checkpoints that monitor the metabolic status of the cell before commitment to phases that expend cellular resources. The utility of the YMC to investigate these processes is the synchrony between metabolic status and transcriptional output with temporal resolution. Additional research will likely uncover other chromatin modifiers that are needed to enact downstream transcriptional regulation of metabolic signaling pathways.

## METHODS

### Yeast strains and metabolic treatment

*Saccharomyces cerevisiae* strains were constructed in the CEN.PK background using standard genetic techniques (Supplementary table 4). Metabolic cycling conditions were performed as previously described previously^3^, except starter cultures were grown in minimal media without sulphuric acid. All strains used were negative for the *petite* phenotype caused by mitochondrial dysfunction, as assessed by the ability to grow on glycerol-containing media. For RNA-seq analysis of the rapamycin-sensitive transcriptome, the BY4741 strain was used. A sample was taken, then rapamycin (30 nM) added and further samples were taken at 45 and 90 minutes. Treatments were performed in duplicate.

### Western blotting

Protein was extracted using an NaOH/TCA extraction method. Standard conditions were used for SDS-PAGE and Western blotting. Antibodies used in this study were: Hxk1 (Novus, NB120-20547), Arp5 (Abcam, ab12099), H3 (Active Motif, 39163), H3K9ac, (Millipore, 06-942), pRps6 (CST, 2211).

### RNA-Seq analysis

RNA was prepared from samples (approximately 1.5 ODs) using the MasterPure^™^ Yeast RNA Purification Kit (Epicentre, MPY03100). The sequencing libraries were prepared from 0.8 μg of RNA/sample using the Illumina TruSeq Stranded mRNA kit (Illumina, 15031047). The quality of the pooled library was checked using the Agilent Bioanalyser 2100 HS DNA assay. The library was sequenced on an Illumina NextSeq Sequencer for the YMC data and on an Illumina HiSeq 2000 platform for the rapamycin dataset. Minimum of 10 million reads per time-point were aligned using Bowtie 2 and analysed using the DESeq2 package^67^. Samples were analysed in duplicate and replicates were combined for final analysis. PCA plots were generated on log-transformed data using the DESeq package^67^. For transcriptional comparative analysis in Figure 2c, time-points with the highest correlation between wild-type and mutant log2-transformed expression within each phase were chosen, which includes the following: wild-type sample 2 vs mutant sample 1 (RC), *r* =0.95; wild-type sample 3 vs mutant sample 3 (OX), *r* =0.91; and wild-type sample 6 vs mutant sample 5 (RB), *r* = 0.96. Functional annotation analysis was performed using DAVID using default parameters^68,69^.

### ATAC-Seq analysis

The ATAC-seq assay and data processing was performed as previously described^70^. ATAC-seq signal in promoters was quantified as the number of fragments mapping to the window 400bp upstream to 100bp downstream from the TSS. For motif analysis, the motifmatchr R package was used to identify motifs from the JASPAR 2016 database^48^ using a p-value threshold of 10^−4^. Accessibility scores for promoters and motif-containing promoters were determined using chromVAR^71^. Nucleosome positions were determined using NucleoATAC^70^, and the first nucleosome at least 50 bp upstream of the TSS was identified as the −1 nucleosome and the first nucleosome downstream of that position as the +1 nucleosome.

### Analysis of DNA content and budding index

Cells were collected from the chemostat and fixed to a final concentration of 70% overnight at 4°C. Cells were washed and resuspended in 50 mM sodium citrate (pH 7.4) before briefly sonicated and treated with RNase A (0.25 mg/ml at 50°C for 2 hours) and. proteinase K (0.75 mg/ml and cells at 50°C for 2 hours). Cells were pelleted and resuspended in citrate buffer containing Sytox Green at 2 μM and DNA content was analysed using a flow cytometer. To assess budding index, cells were counted using a haemocytometer. Budding percentage was calculated as number budded/total cells (minimum of 400 counted per time-point).

### Data availability

RNA-seq and ATAC-seq data are available under the NCBI accession GSE101290.

## Acknowledgements

This work was funded by National Institutes of Health R35GM119580 to AJM. We wish to thank the Department of Genetics Bootcamp for technical assistance with the ATAC-seq sample processing.

## Author contributions

AJM and GJG conceived study and interpreted results. GJG planned experiments, performed experimental analysis and RNA-seq computational analysis. ANS performed ATAC-seq experiments and computational analysis. WJG supervised ATAC-seq analysis. KMW performed additional RNA-seq and ATAC-seq analysis. DAK established the YMC bioreactor system. GJG, AJM, and DAK wrote the manuscript. All authors approved the manuscript.

## Materials & Correspondence

Correspondence should be addressed to AJM.

---

**Supplementary Figure 1.**
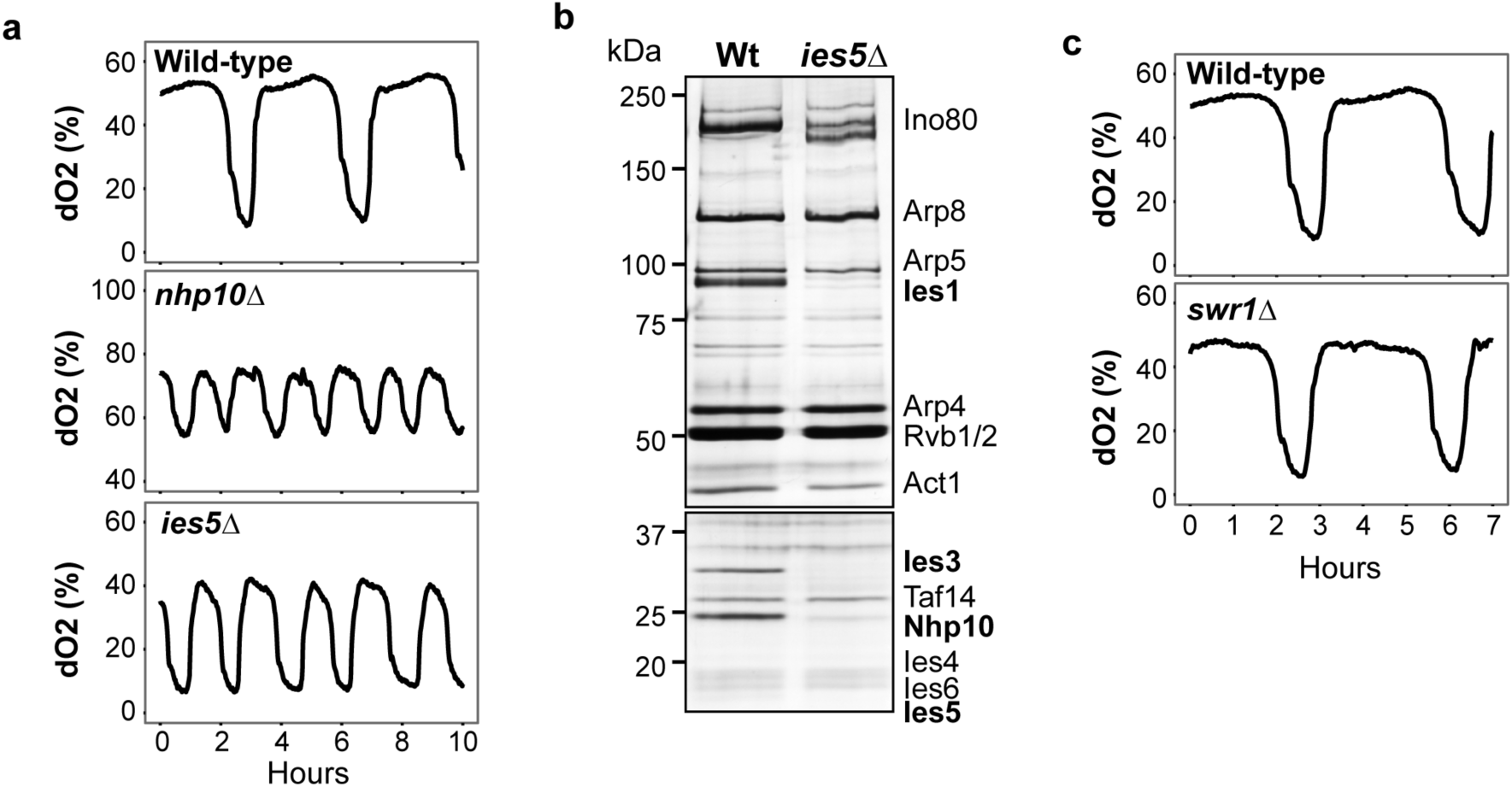
The Nhp10 module of the INO80 complex, but not Swr1, is needed for YMC maintenance. **(a)** Deletion of *NHP10* or *IES5* caused similar YMC defects. YMC dissolved oxygen (dO2) traces in wild-type and the indicated mutants are shown. **(b)** Deletion of *IES5* reduces the levels of Ies1, Ies3 and Nhp10 associated with the INO80 complex. Shown are silver stained SDS-PAGE gels of Ino80-FLAG purified from either wild-type (Wt) or *ies5*Δ strains. Molecular weight is shown as kilodaltons (kDa) on left. Previously identified subunits^26^ are shown on right. **(c)** Deletion of *SWR1* had no effect on the YMC. YMC dissolved oxygen (dO2) traces in wild-type and the *swr1*Δ mutant are shown.

**Supplementary Figure 2.**
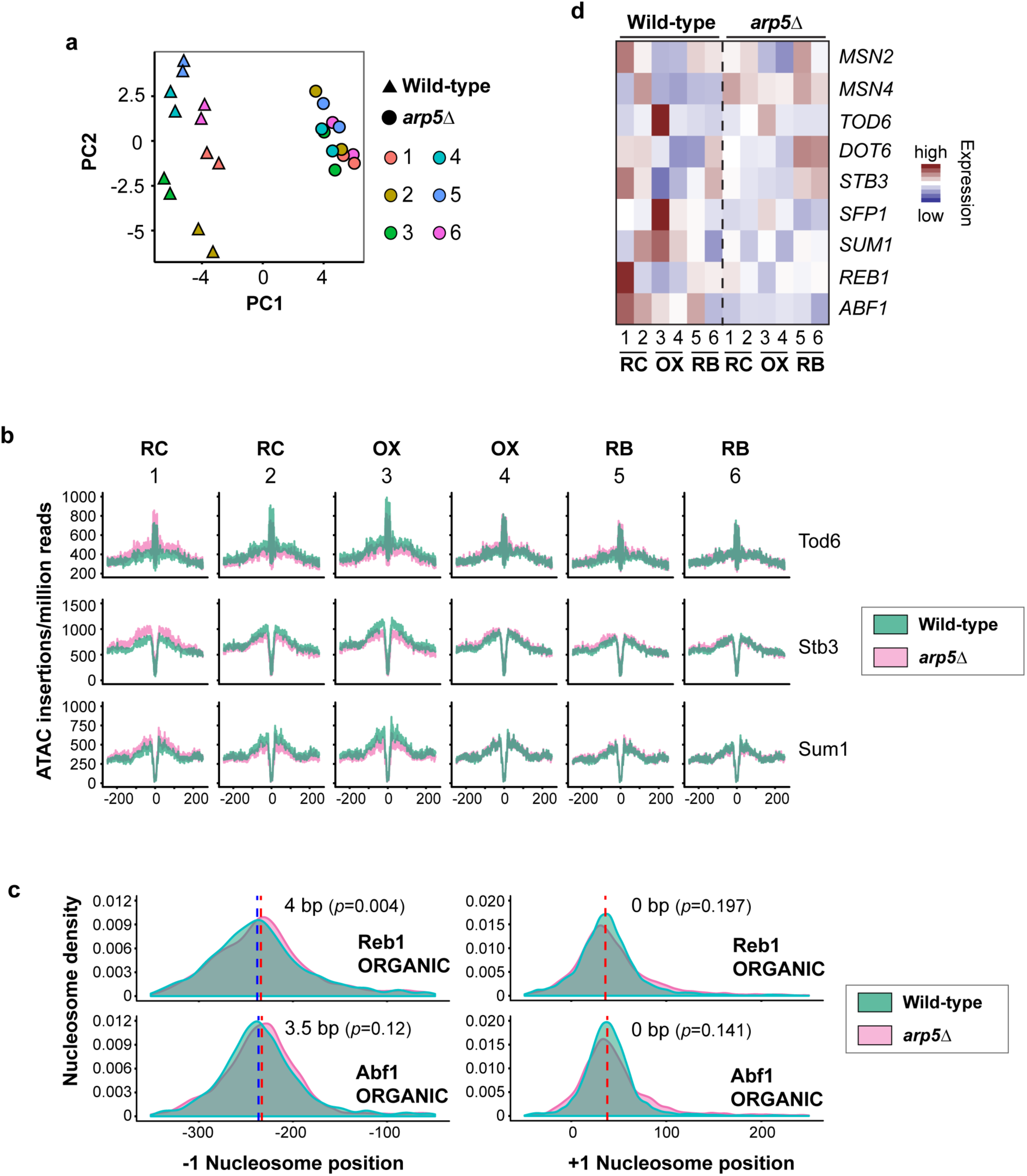
INO80 regulates chromatin accessibility at several metabolic gene promoters. **(a)** Principal component analysis (PCA) plot of ATAC-seq data for wild-type and *arp5*Δ YMC samples taken at timepoints shown in Fig. 2a. PC1 and PC2 are shown. Technical replicates are displayed for each time point and strain. **(b)** ATAC accessibility is shown as the number of Tn5 adaptor insertions per million reads around the indicated transcription factor motif for wild-type (green) and *arp5*Δ (pink) samples. **(c)** +1 and −1 nucleosome densities, relative to transcription start sites containing Abf1 or Reb1 motifs, were obtained from ORGANIC profiles^49^and identified as previously described using NucleoATAC^70^. Wild-type (green) and *arp5*Δ (pink) samples across all time-points of the YMC are shown. Red/blue dotted lines denote the median nucleosome position for wild-type and *arp5*Δ respectively. p = 0.01188 (Abf1) and 0.00408 (Reb1) based on permutation test and adjusting for multiple hypothesis testing. **(d)** Heatmaps showing expression of genes with the indicated transcription factor motif in the promoter. Mean expression is shown as Fragments Per Kilobase of transcript per Million mapped reads (FPKM).

**Supplementary Figure 3.**
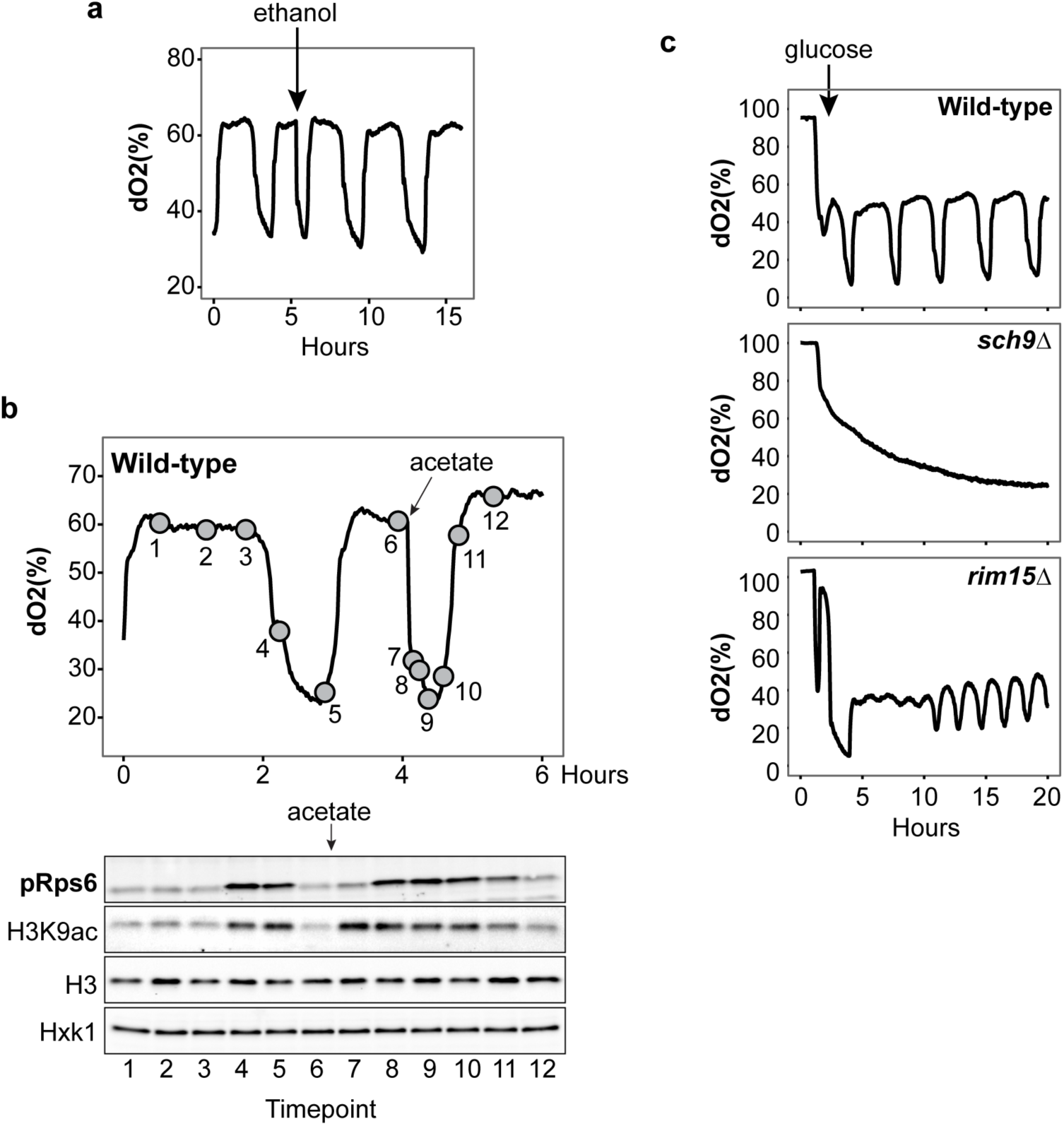
TORC1 signaling regulates the YMC. **(a)** Control for Fig. 6d in which rapamycin was dissolved in ethanol. Shown is the YMC dissolved oxygen (dO2) trace where 10 μl of ethanol is added. Following induction into the oxidative phase, the YMC progresses normally. **(b)** Acetate treatment induces oxidative phase and increases TORC1 activity. Samples were taken at indicated timepoints across the wild-type YMC before and after treatment with sodium acetate (1 mM). Samples were used for western analysis with an antibody that recognizes the phosphorylated TORC1 substrate, mammalian homologue ribosomal protein S6 (Rps6). Also shown is acetylated histone lysine 9 (H3K9ac). Histone H3 (H3) and hexokinase 1 (Hxk1) are shown as a loading controls. **(c)** Deletion of *SCH9* or *RIM15* causes YMC defects. YMC dissolved oxygen (dO2) traces in wild-type and the indicated mutants are shown.

